# The precision of signals encoding active self-movement

**DOI:** 10.1101/2023.09.20.558633

**Authors:** Joshua D. Haynes, Maria Gallagher, John F. Culling, Tom C.A. Freeman

## Abstract

Everyday actions like moving the head, walking around and reaching out to grasp objects are typically self-controlled. This presents a problem when studying the signals encoding such actions because active self-movement is difficult to experimentally control. Available techniques demand repeatable trials, but each action is unique, making it difficult to measure fundamental properties like psychophysical thresholds. Here, we present a novel paradigm that can be used to recover both precision and bias of self-movement signals with minimal constraint on the participant. The paradigm takes care of a hidden source of external noise not previously accounted for in techniques that link display motion to self-movement in real time (e.g. virtual reality). We use head rotations as an example of self-movement, and show that the precision of the signals encoding head movement depends on whether they are being used to judge visual motion or auditory motion. We find perceived motion is slowed during head movement in both cases, indicating that the ‘non-image’ signals encoding active head rotation (motor commands, proprioception and vestibular cues) are biased to lower speeds and/or displacements. In a second experiment, we trained participants to rotate their heads at different rates and found that the precision of the head rotation signal rises proportionally with head speed (Weber’s Law). We discuss the findings in terms of the different motion cues used by vision and hearing, and the implications they have for Bayesian models of motion perception.

**NEW AND NOTEWORTHY:** We present a psychophysical technique for measuring the precision of signals encoding active self-movements. Using head movements, we show that: (1) precision declines when active head rotation is combined with auditory as opposed to visual motion; (2) precision rises with head speed (Weber’s Law); (3) perceived speed is lower during head movement. The findings may reflect the steps needed to convert different cues into common currencies, and challenge standard Bayesian models of motion perception.

## INTRODUCTION

Bodily movement is a key part of everyday life. Our eyes, head, limbs and torso are seldom at rest. Action therefore sets the backdrop in which perceptual systems normally operate, with many everyday tasks relying on information about current self-movement. This is derived from a number of perceptual signals, some based on images such as retinal flow, and some based on non-image sources including the vestibular and motor systems.Information from non-image sources also plays a role in interpreting images, allowing the observer to differentiate between self-generated movement and movements of external objects.

Success in these active tasks is constrained by two fundamental types of error, namely the precision and accuracy of the underlying perceptual signals. Precision is driven by internal and external noise and corresponds to the width of the distribution of the underlying perceptual signal as it varies across time. Accuracy, on the other hand, corresponds to the distribution’s average and is usually referred to as bias. While a lot is known about the precision and accuracy of image signals, especially in vision and hearing, much less is known about the errors accompanying non-image signals. This is especially the case when the self-movement is ‘active’ (i.e., self-controlled), partly because it is difficult to apply standard psychophysical techniques to actions that are under participant control. We therefore developed a new way to measure precision in these circumstances, using head rotation as an example of self-movement. The technique makes no attempt to differentiate between the various sources of non-image information that are used to encode active self-movement. Rather, it assumes that they are grouped together to provide a single non-image signal, and it is the precision of this composite signal we measure. Our technique relies on using image signals as a comparison, which allowed us to compare the results when using vision or hearing in Experiment 1. We then used the same technique in Experiment 2 to investigate whether non-image precision obeys Weber’s law, that is whether precision scales with magnitude (e.g. Altman & Viskov, 1977; De Bruyn & Orban, 1988; Mallery et al., 2010). Investigating Weber’s law also allowed us to compare the precisions of image and non-image signals.

In a typical precision-measuring task, participants are asked to compare a fixed standard stimulus with a range of test stimuli shown over a series of trials. The values assigned to the test are usually controlled by a method of constant stimuli or a staircase procedure, both of which provide a measure of a just-noticeable-difference that can be used to estimate the precision of the underlying signal. Crucially, these methods rely on the ability to repeat a set of stimuli over trials. Self-controlled self-movements, however, are not repeatable: every instance of every action is unique. To measure non-image signal precision, this variability must be accounted for, both within and across trials.

One solution is to use ‘passive’ self-movement because the action is then controlled by the experimenter. The best-known examples come from vestibular research, where participants are moved on a chair or platform (Brooks & Cullen, 2019). Notable examples also come from studies of perceived stability during eye movement, where various contraptions and implements have been used to passively rotate the eye (Merton, 1964; Skavenski, 1972).

But active non-image signals also include efferent sources, such as copies of motor commands (Von Holst, 1954), not just the vestibular, somatosensory and, in the case of passive rotation of the eye, the proprioceptive cues that passive stimulation generates (Cullen & Zobeiri, 2021; Israel & Warren, 2005; St George & Fitzpatrick, 2011; Tuthill & Azim, 2018). Passive stimulation therefore reduces the number of non-image sources down to one or two, potentially producing cue conflicts with those that have been silenced.

Instead, we focus on active self-movement, specifically head rotation, where non-image sources consist of vestibular cues, motor commands, and proprioceptive feedback, plus any number of somatosensory cues, such as the gliding of hair across the back of the neck. Our method for measuring non-image signal precision uses two experimental phases, combined with a novel analysis that accounts for the variability of self-movement across trials. The paradigm is sketched in Figure 1. Based on two-interval forced-choice, the participant makes self-controlled back and forth head rotations in the first interval of each trial of Phase 1, and an auditory or visual stimulus appears in the 3^rd^ sweep (see bottom left panel). This stimulus is head-centred – it moves with the participant – and is used to mark which portion of the head movement to judge. We refer to this interval as the ‘standard’. In the second ‘test’ interval of Phase 1 shown in the 2^nd^ column of the figure, an auditory or visual stimulus is again shown, but this time with the head stationary. The stimuli move with a trajectory defined by the recorded head movement but scaled up or down by a multiplicative factor we call ‘motion gain’. Hence, the pattern and duration of motion experienced in the two intervals is the same, apart from overall magnitude, and is encoded by different signals: non-image in the first, and image-based in the second. Participants indicate which interval appears to ‘move more’. We avoided the terminology ‘faster’ or ‘further’ because cue preference depends on modality: for vision, participants prefer speed rather than displacement and duration (Freeman et al., 2018; Reisbeck & Gegenfurtner, 1999), whereas for hearing the reverse is true (Carlile & Best, 2002; Freeman et al., 2014). Motion gain is manipulated across trials using a Method of Constant Stimuli, resulting in a psychometric function that includes two sources of internal noise, one based on the image signal (e.g. visual or auditory motion) and one based on the non-image signal. To tease these two sources apart, Phase 2 isolates the internal noise of the image signal, using the same set of head movement recordings from Phase 1 to move the stimuli in the same trial-by-trial order, but with the head always stationary. Again, the second interval is a scaled version of the first. The precision of the non-image head rotation signal can then be recovered from Phase 1, with image precision now known.

**Figure 1.**
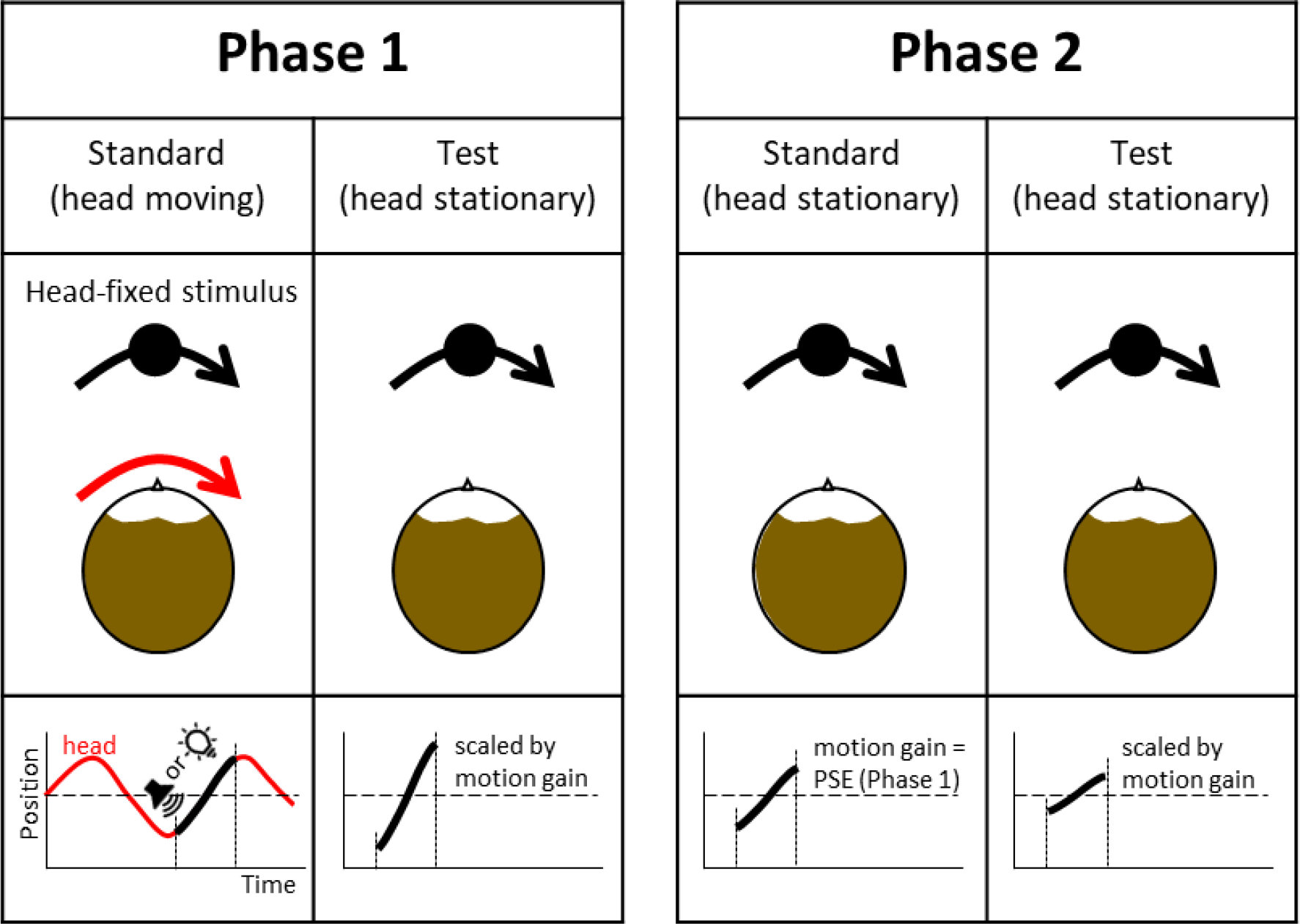
Schematic of the two-phase procedure for measuring the precision of non-image signals encoding active self-movement. We use head rotation as an example. Phase 1 consists of two intervals: a standard interval, in which the head moves and a head-fixed stimulus (visual or auditory) appears ‘on the nose’ in the 3rd sweep, and a test interval, in which the same movement of the stimulus, scaled by the motion gain, is replayed but with head stationary. Phase 2 also consists of two intervals, both with the head stationary. Here, the motion gain used to scale stimulus motion in the standard interval is set to the Point of Subjective Equality found in Phase 1. For both phases, a Method of Constant stimuli is used to manipulate motion gain across trials and construct psychometric functions. These are then used to determine the precision of the image signal in the head-stationary intervals, and the non-image signal in the head-moving interval, based on a model described in the Appendix.

In Experiment 1, we used this novel technique to measure non-image precision accompanying head rotation, using either auditory or visual stimuli. We predicted that the modality used to deliver image motion should not affect the precision of the non-image signal measured. In Experiment 2, we used auditory stimuli to investigate whether the precision of the non-image signal obeys Weber’s law (i.e., scales with magnitude). Note that Phase 1 of the paradigm also provides information about bias, specifically whether the magnitude of perceived motion is the same with or without head movement. This is interesting in its own right, partly because it is well known that objects pursued by an eye movement appear slower (Aubert, 1887; Fleischl, 1882). Analogous perceptual slowing has been demonstrated for passive stimulation of the vestibular system (Garzorz et al., 2018) and active touch (Moscatelli et al., 2019), and has also been implicated for the auditory system during head rotation (Freeman et al., 2017). But as far as we are aware, whether head rotation produces the same slowing is currently not known for either vision or hearing.

## METHODS

### Experiment 1

#### Stimuli & Materials

Auditory stimuli were played over a 2.4m diameter ring of 48 Cambridge Audio Minx speakers as shown in Figure 2. The room was sound treated and completely dark. The speakers were controlled by two MoTU sound cards, each linked to four amplifiers. Intensity was normalised across individual speakers. The stimuli consisted of white noise spatially windowed by a Gaussian distribution (σ = 5.25° in power, equivalent to 0.7 of the speaker spacing i.e., σ = 7.5° in amplitude). We have previously shown that this value avoids aliasing artifacts in our speaker system that could occur if the Gaussian distribution is undersampled, while at the same time avoiding the sound becoming too diffuse (Stevenson-Hoare et al., 2022). The noise was sampled at a rate of 48 KHz with a peak level of 70 dB. The position of the spatial Gaussian was refreshed at a rate of 240 Hz, a rate set by the motion tracker described below. The result was a ‘blob’ of noise that could be moved smoothly across the speakers. The actual motion path taken was determined by the measured head movements, using the motion gain parameter to scale its magnitude.

**Figure 2.**
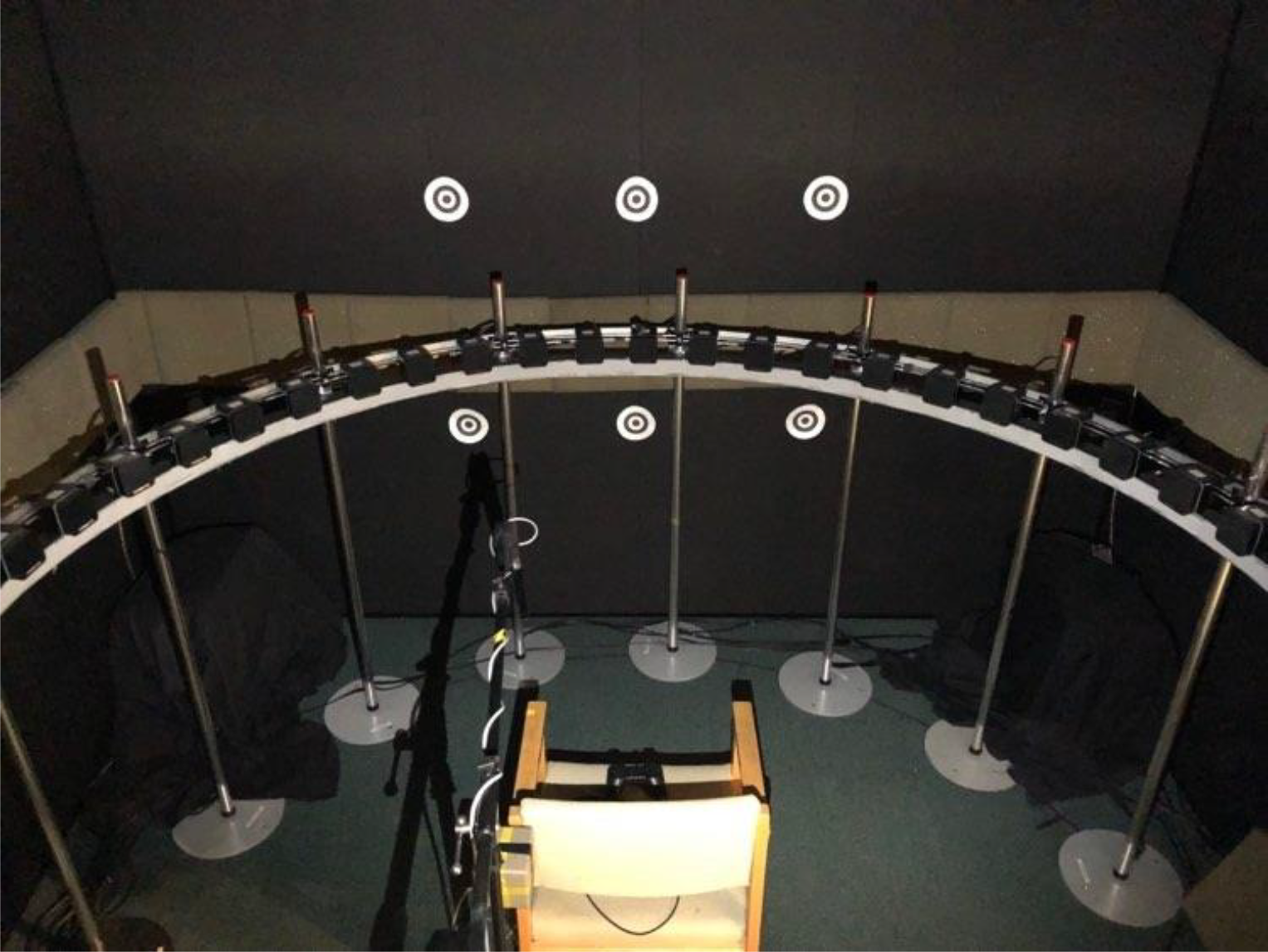
Laboratory set-up.

Visual stimuli were presented to the participant using an AdaFruit NeoPixel strip of 342 LEDs driven by a single Arduino Uno microcontroller. The LED strip was positioned just below the speakers, as shown in Figure 2, and was driven at a framerate of 40 Hz. The strip subtended 128° either side of straight ahead. This yielded an LED spacing of 0.75°. To ensure that the LEDs presented stimuli at a comfortable brightness, a single layer of 1.2f neutral density filter reduced the intensity of the display. As with the auditory stimuli, smoothly moving stimuli were created by using a Gaussian distribution that spatially windowed the LED output for each display frame (σ = 1.05°). In order to prevent individual LEDs being visually resolved, the strip was placed in a curved enclosure with one open side that was covered by three layers of diffuser gel at a distance of 35 mm, blurring the image. The overall size of the resulting blob was increased slightly by the diffuser (σ = 1.07°), which we confirmed using a Minolta LS100 photometer and an array of small apertures. The peak luminance of the blob was ∼0.042 cd/m^2^.

#### Head Tracking

Head movement was measured using a Polhemus Liberty tracker that sampled position at a rate of 240 Hz. The tracker was mounted to a head band worn by the participant. For the head-moving interval of Phase 1, the head-tracking data were used to detect the 3^rd^ sweep in real-time and keep the subsequent auditory or visual stimulus head-centred (i.e., motion gain = 1). To detect a change in head-movement direction, we convolved the head tracker samples with a finite difference filter to obtain a smoothed derivative. The filter was 13 samples long, meaning there was a 7-frame delay in detecting the head-turn (∼30msec). An example waveform is shown in Figure 3A, with the detected 3^rd^ sweep shown in black and blue.

**Figure 3.**
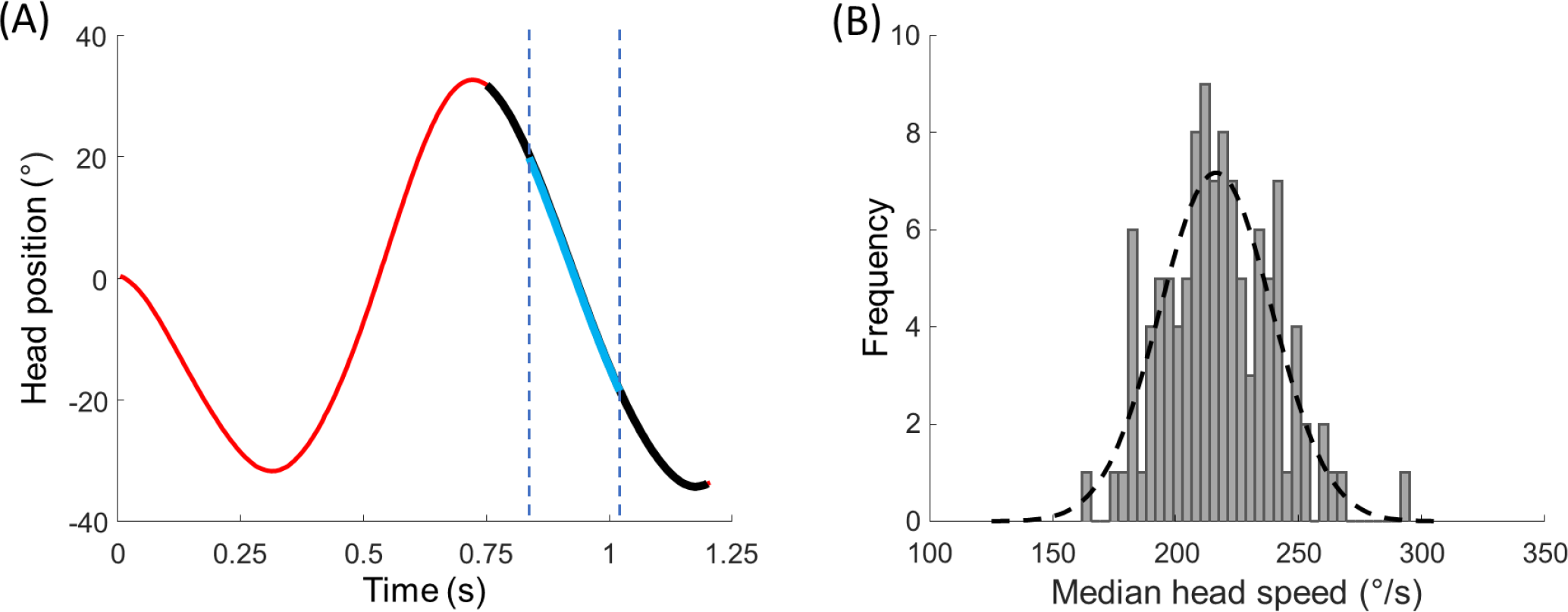
(A) Example head movement waveform. The black portion corresponds to the 3rd sweep as detected by the algorithm described in the text. The visual or auditory stimulus appeared during this time. The blue portion defines the region of interest over which median head speed was calculated for analysis. (B) An example distribution of median head speeds for a single repetition of Phase 1 (110 trials) for one participant. The dotted line shows the best fitting Gaussian, which was used to determine the mean and standard deviation of the distribution.

#### Procedure

In Phase 1, each trial consisted of a ‘head-moving’ standard followed by a ‘head-stationary’ test. The start of the first interval was signalled by a short beep (0.25s) followed by momentary pause to check the head was centred before the experiment moved on. ‘Centred’ was defined as 10 consecutive head-tracker samples within ±7.5° of the centre of the LED/speaker array. Participants then began to rotate their head smoothly left and right or vice versa. While it was their free choice, we found some participants alternated the start direction from trial to trial, while others mostly started in the same direction. The auditory or visual stimulus appeared during the 3^rd^ sweep and moved with the head. The start of the head-stationary test interval was signalled using a blue blob that appeared for 0.2s, with progress again paused to check the head was centred. Note that the motion in the test interval was based on the 3^rd^ sweep recorded in the standard interval only: the dead-time created by the initial 2 sweeps was skipped. In Phase 2, the same beep and light were used to identify the start of each interval, with both intervals head-stationary and the initial 2 sweeps skipped. Each replication of Phase 1 and Phase 2 contained the same number of trials, based on the same head movement recordings, shown in the same order.

Psychometric functions were collected based on a Method of Constant Stimuli using 11 motion gain values. For Phase 1, these ranged from 0.2 – 1.2 in 0.1 steps for the auditory condition, and 0.3 – 1.1 in 0.06 steps for the visual condition. The ranges were based on pilot experiments that showed the visual condition produced steeper psychometric functions than the auditory condition. Each motion gain was repeated 10 times, yielding 110 trials per session. For Phase 2, the same step sizes were used, but the range was centred on the Point of Subjective Equality (PSE) calculated from Phase 1. This ensured that the precision of the image-motion signal we estimated for each replication of Phase 1 and 2 were based on motion gains centred on a comparable value. The PSE was derived using the Palamedes toolbox (note the model described in the Appendix returns the same PSE as the toolbox).

Participants sat in the centre of the speaker/LED ring and wore the head tracking equipment (in Experiment 2 they also wore an eye tracker). Head position was checked with a laser crosshair mounted above the centre of the ring, which enabled the interaural axis and speaker ring to be aligned. The head tracker was boresighted with the participant facing forwards and pointing their head towards the central speaker. Boresighting was repeated at the start of each replication of each phase of the experiment.

Each participant repeated three pairs of Phases 1 and 2 for each modality. Three out of five participants carried out the auditory condition first.

#### Head movement analysis

To analyse the head movements after data collection, position samples were first smoothed using MatLab’s ‘lowpass’ function with a passband of 8 Hz. The temporal derivative was then taken and the median velocity calculated over a portion of the 3^rd^ sweep that ranged from 20-60% of the sweep length (shown in blue in the example waveform of Figure 3A). This Region-Of-Interest (ROI) was adopted because it maximised the number of head-movement samples and goodness-of-fit of the psychometric function (see Appendix for evaluation). Figure 3B shows an example for one participant of the distribution of these median velocities for one run of Phase 1. For modelling purposes, the distribution of each 110-trial run was fit with a Gaussian (dotted line) to extract a mean and standard deviation.

#### Psychophysical analysis

The distribution shown in Figure 3B emphasises the fact that, as with other self-movements, head rotation varies across time. Using motion gain therefore seems a good way of controlling for this variability because it links the related image motion to the ongoing self-movement in real-time. The patterns of motion are therefore identical, meaning the only difference between signal inputs is speed and displacement – duration is fixed. On the face of it, therefore, motion gain provides the experimenter with a repeatable parameter that can be used to define a psychometric function or drive a staircase. Examples are provided by Serafin et al (2013) and Steinicke et al (2009), who plot psychometric functions defined by changes in motion gain within visual and acoustic virtual reality set-ups, respectively. However, closer inspection of their figures suggests a consistent feature not accounted for by fitting a standard cumulative Gaussian: on occasions, their data appear to asymptote more than a constrained lapse rate parameter would allow (e.g. <6%, as suggested by Wichman & Hill, 2001). In the Appendix, we construct a model of the psychophysical task that shows why. The model emphasises that motion gain is not always a good shorthand for the actual stimulation experienced by the participant, namely the magnitude of motion (speed or displacement).

In keeping with standard signal detection theory, the model assumes that participants base their judgement on a point estimate of stimulus magnitude (e.g., the peak speed of head and image movement, or average speed, or displacement). Crucially, the point estimates vary across trials due to the external noise introduced by variable self-movement, as well the internal noise. The external noise produces some surprising effects (see Figure A1). First, the function’s true slope is steeper than the best-fitting single cumulative Gaussian. Second, as the variability of the self-movement increases, the function’s asymptotes depart markedly from 0 and 100%, much further than a typical constrained lapse rate parameter of 6% would allow.

Following standard practice, we assume that internal and external noise is Gaussian distributed. The precision of a given signal is therefore defined by its standard deviation. If the self-movement did not vary at all, the precision of the non-image signal could be calculated by standard fitting of a cumulative Gaussian to the psychophysical data, and then applying the ‘variances sum’ law to both phases. Thus, for Phase 1, 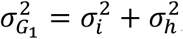, where the subscripts correspond to the cumulative Gaussian fit to the data, the image signal (auditory or visual), and the non-image signal encoding head rotation. For Phase 2, 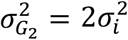 hence the precision of the non-image signal 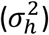 in Phase 1 can be found by substitution. But when self-movement varies, this standard approach is an approximation at best. Variable self-movement adds external noise that varies across the psychometric function because it is scaled by motion gain; hence the assumption of a single cumulative Gaussian is not correct. We develop the appropriate formulae in the Appendix and show how these can be used to extract the internal noise of the image signal 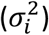 and non-image signal 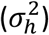 from the two phases of our experiment. Similar formulae can be applied to a more typical motion gain scenario used in virtual reality set-ups, where both self-movement and image motion are shown at the same time (e.g., Cherni et al., 2020; Nilsson et al., 2018; Steinicke et al., 2009; Serafin et al).

### Experiment 2

#### Stimuli & Procedure

The aim of Experiment 2 was to determine signal precision as a function of head and stimulus speed. Before each replication of the main experiment, a training session was therefore run to help participants rotate their heads at one of the five target speeds. The training stimuli were audiovisual, consisting of visual and auditory blobs as used in Experiment 1. These moved in synchrony. During the main experiment, however, only auditory stimuli were used. The procedure for the main experiment was the same as Experiment 1, consisting of two experimental phases linked by the set of head movements recorded in the first. Each trained head speed was investigated by completing training and main experiment pairs. This process was repeated three times, yielding 15 training and main experiment pairs presented in a random order.

#### Head Speed Training Sessions

Training sessions consisted of a two-stage process that was run ahead of each replication of the main experiment. In stage 1, participants were asked to track an audiovisual stimulus with their head. The stimulus moved independently along a sinusoidal path at a frequency of 1Hz at one of five amplitudes: 5°, 12.5°, 20°, 27.5°, 35°. These correspond to median target speeds from 30.0°/s to 209°/s for the ROI defined in Experiment 1. Five and three-quarter periods were shown for each speed to generate 12 sweeps. In stage 2, participants attempted to reproduce the trained head speed, this time using a head-stationary audiovisual fixation target moving with the nose as a guide (motion gain = 1). Again, they completed five and three-quarter periods, determined by recording the number of head direction reversals detected in the head tracking as described in Experiment 1. The accuracy of head rotation was assessed by calculating the median head speed for each of the final 10 head sweeps as described in Experiment 1. If 7/10 sweeps had a median within 5°/s ±5% of the desired training speed, performance on that training run was deemed sufficiently accurate. If not, the participant was given feedback on how many sweeps were accurate, and how many were too fast and/or slow, and the run repeated. Participants had to complete at least three training runs, with at least one successful run before progressing to each replication of the main experiment.

#### Eye Tracking and Analysis

Eye movements were tracked using a Pupil Labs Pupil Core head mounted eye tracker. The tracker had a 120 Hz sampling frequency and a front-facing world camera. The camera was used for calibration by having participants look at a 3 by 2 array of calibration points that can be seen in Figure 2. These were used to convert the eye tracker’s normalised units into degrees. To analyse, the data were first drift corrected trial by trial. To calculate fixation accuracy of the head-stabilised stimulus, a spatial ROI of ±3° from the centre of the auditory blob was defined and then the percentage of samples measured within it was calculated for the 3^rd^ sweep. For reference, we also calculated an equivalent measure corresponding to a participant perfectly counter-rotating their eyes to maintain fixation on a world-stationary target.

#### Participants

All observers gave informed consent, and the experimental procedures were approved by the School of Psychology, Cardiff University Ethics Committee (EC.12.04.03.3123GRA2). In Experiment 1, five participants took part in the experiment (2 female, 3 male). Two participants were naïve to the purposes of the experiment and three were experimenters. Participants wore spectacle correction if required. In Experiment 2, three experimenters and seven participants studying psychology at Cardiff University took part in the experiment (2 male, 8 female). The latter group were unaware of the aims of the experiment. Participants completed at least two replications of the experiment, with eight participants completing three. Eye movements were recorded for nine of the participants. Only one of these normally wore spectacles, which were removed to allow the eye tracker to operate.

The code used for fitting the model using the two-phase paradigm, together with the raw data and summary data, can be found here (https://osf.io/qcz7w/?view_only=2e2bb846820d4862bcb02a036d3ee815)

## RESULTS

### Experiment 1: Precision of signals encoding active head rotation

Figure 4A shows mean the mean precision of the non-image head-rotation signal (bars) across the five participants together with their individual data (solid points). Non-image signals were less precise in the auditory condition, producing an increase in the standard deviation of the underlying signal distribution as defined by the model described in the Appendix (t(4) = 4.19, p = .01). Contrary to our prediction, therefore, modality appears to matter. Figure 4B shows that head speeds were slightly faster when an auditory target appeared in the 3^rd^ head sweep compared to a visual one, however this difference was non-significant (t(4) = 1.76, p = .15). It is unlikely, therefore, that the marked difference in non-image signal precision measured using visual and auditory stimuli could be explained by a scaling of precision with magnitude (i.e. Weber’s law). This point is explored further in Experiment 2, where Weber’s Law was investigated more thoroughly.

**Figure 4.**
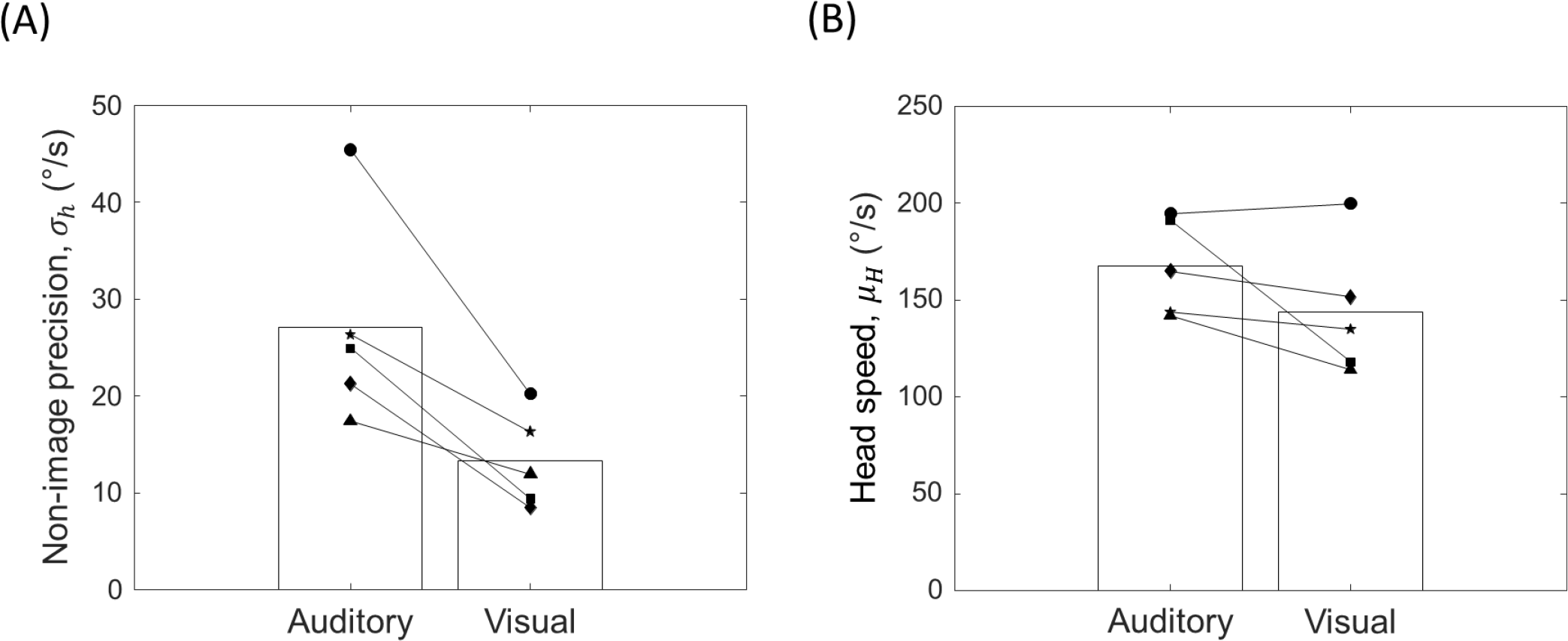
(A) Precision of the non-image signal encoding head rotation for the two stimulus conditions. Precision is defined as the standard deviation of the underlying signal distribution, as defined by the model in the Appendix. Note therefore that larger standard deviations correspond the less precise signals. Bars correspond to the mean of the individual data points shown as solid symbols. (B) Head speed using the same format.

Figure 5 shows the mean PSEs obtained from Phase 1. The PSEs are remarkably similar for vision and hearing (t(4) = -0.71, p = .52). The PSEs are around 0.7, meaning that stimuli had to be slowed by 30% when they moved passed a stationary participant, in order to be perceived as the same speed as the head moving interval. For vision, this finding resembles the Aubert-Fleischl phenomenon (Aubert, 1887; Fleischl, 1882), where motion appears slower if stimuli are followed by a smooth eye pursuit. Our data show that perceived slowing occurs for head movement too, albeit for auditory and visual stimuli that are linked directly to the self-movement as opposed to being pursued in a more typical closed-loop manner.

**Figure 5.**
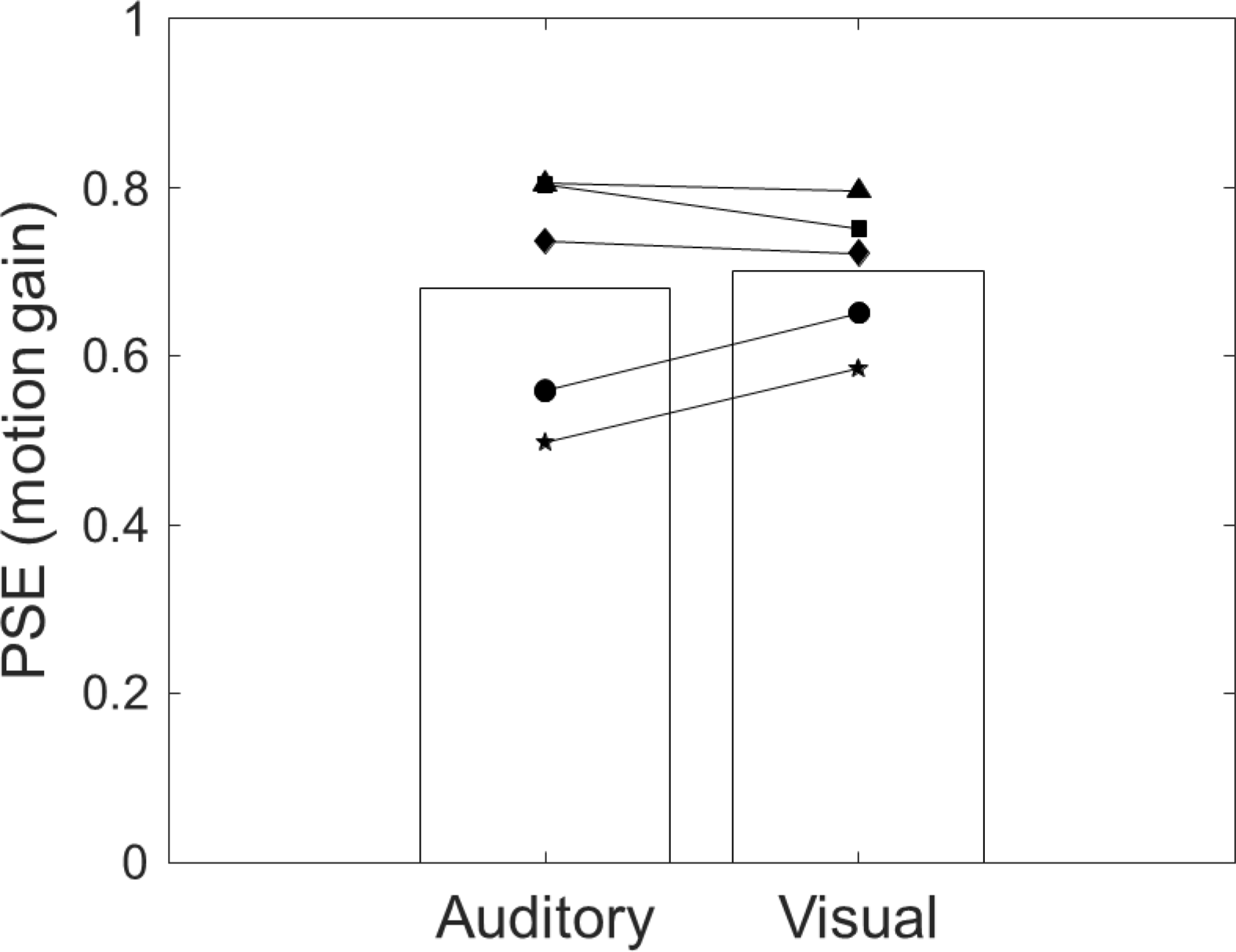
PSEs for the psychometric functions collected in Phase 1. These indicate the relative bias between image and non-image signals for both modalities. A value less than one means that head tracked stimuli appear slower.

### Experiment 2: Do non-image signals obey Weber’s law during active head rotation?

For many sensory systems, discrimination thresholds scale proportionally with stimulus magnitude over a wide range (Weber’s Law: Laming, 2009). In Experiment 2, we investigated the extent to which non-image signals accompanying active head rotation adhered to Weber’s law by training participants to rotate their heads at different speeds. The same two-phase protocol described in Experiment 1 was used to measure non-image precision at each trained speed, using auditory stimuli only. These data also allowed a more direct comparison between non-image and image precision. That was not possible in Experiment 1 because the two-phase protocol forces the motion gain of the standard in Phase 1 to be different from that used in Phase 2 (i.e., gain = 1 and PSE, respectively). By manipulating head speed in Experiment 2, signal precision can be described as a function of magnitude, allowing the comparison to be made.

Figure 6A plots the mean head speed participants executed in the main experiment. The dotted line indicates the speeds targeted in the training runs. The head movement was reasonably accurate in the main experiment, producing a well separated set of rotation speeds that covered a wide range (F(4,45) = 16.89, p < .001).

**Figure 6.**
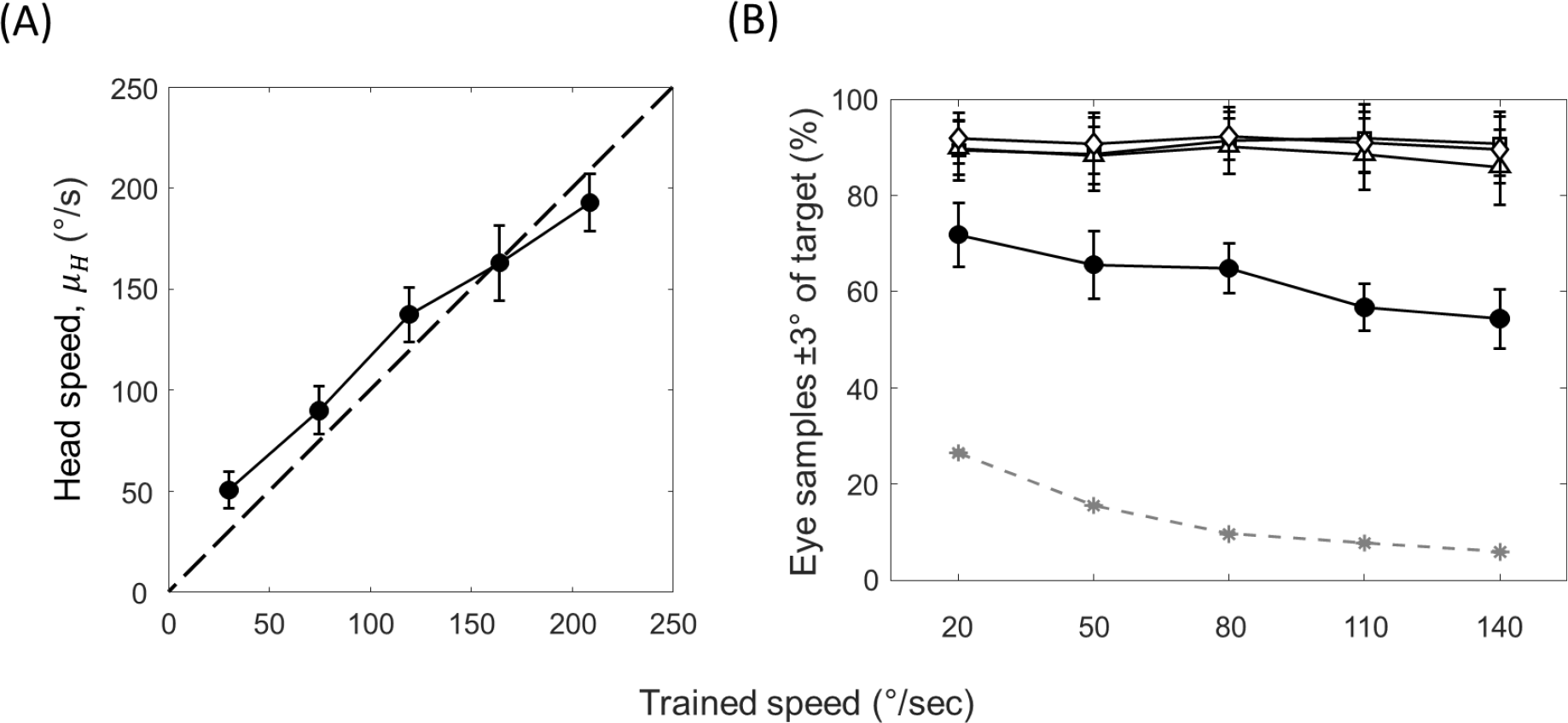
(A) Head velocity over ROI defined in Figure 3A. The dashed line shows the actual trained speed. (B) Eye fixation accuracy expressed as % of samples ±3° of head-centred target. Open symbols correspond to the three head-stationary intervals (one in Phase 1, two in Phase 2) and closed circles the head-moving interval. The stars and dashed line correspond to the equivalent measure if participants had perfectly counterrotated the eye to remain fixed in world-centred coordinates. Error bars are ±1SE.

Figure 6B plots the percentage of eye position samples within a spatial ROI of ±3°, as a function of trained head speed. As can be seen, all head-stationary intervals (open points) produced a similar level of performance. Accuracy was lower for the head-moving interval but was still quite high. As a reference, the dashed line plots the same measure assuming participants had perfectly stabilised eye position via counter-rotation so as to fixate a world-stationary point.

Figure 7A plots the non-image signal precision (filled circles) and auditory signal precision (open circles) as a function of the mean head or stimulus speed, respectively. The latter compresses horizontally because the speeds are set by the PSE obtained from Phase 2 of the main experiment. This corresponds to a motion gain of around 0.7 (see Figure 8 for the PSEs at each training speed). The horizontal compression is therefore around 30% compared to the closed circles.

**Figure 7.**
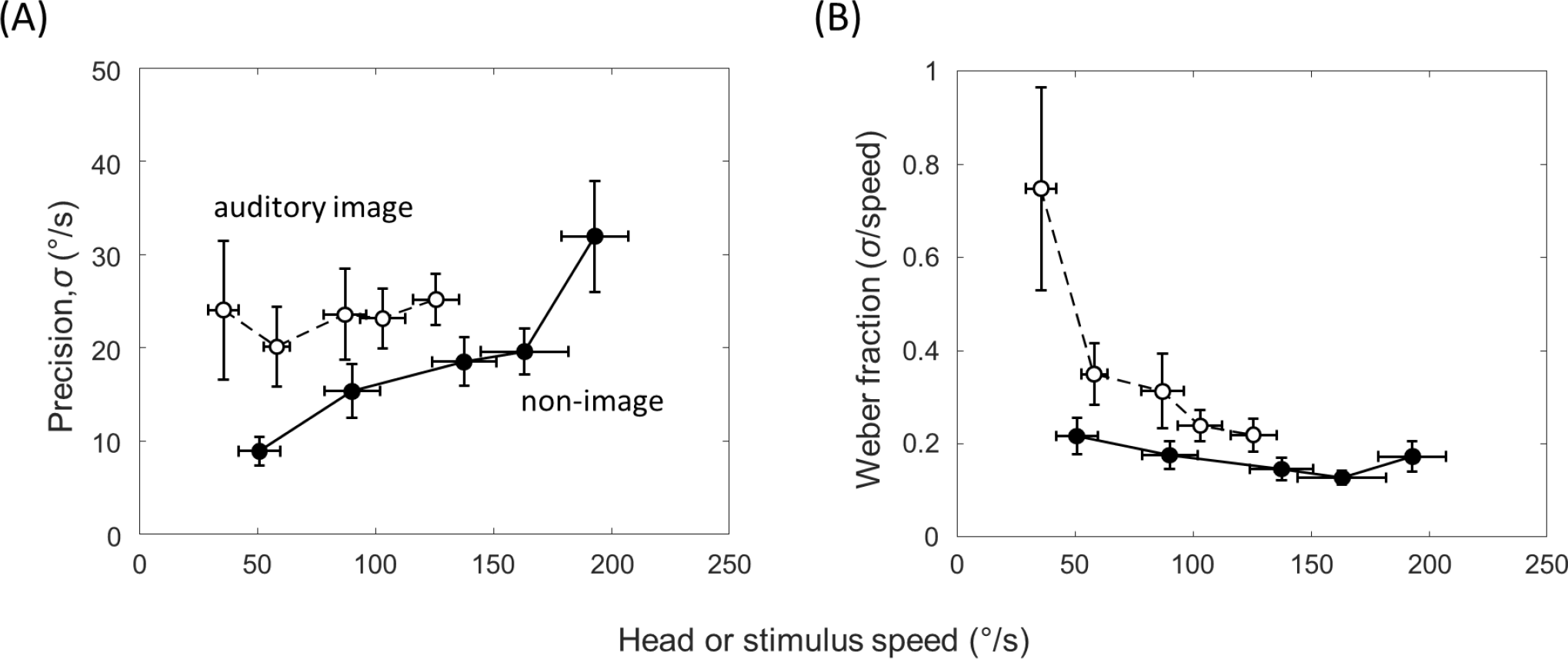
(A) Precision as a function of stimulus speed for the auditory image signal (open symbols) and non-image signals (closed symbols). Precision is defined as the standard deviation of the underlying signal distribution defined by the model in the Appendix. Note therefore that larger standard deviations correspond the less precise signals. (B) The same data expressed as Weber fractions i.e. standard deviation / speed. Error bars are ±1SE.

**Figure 8.**
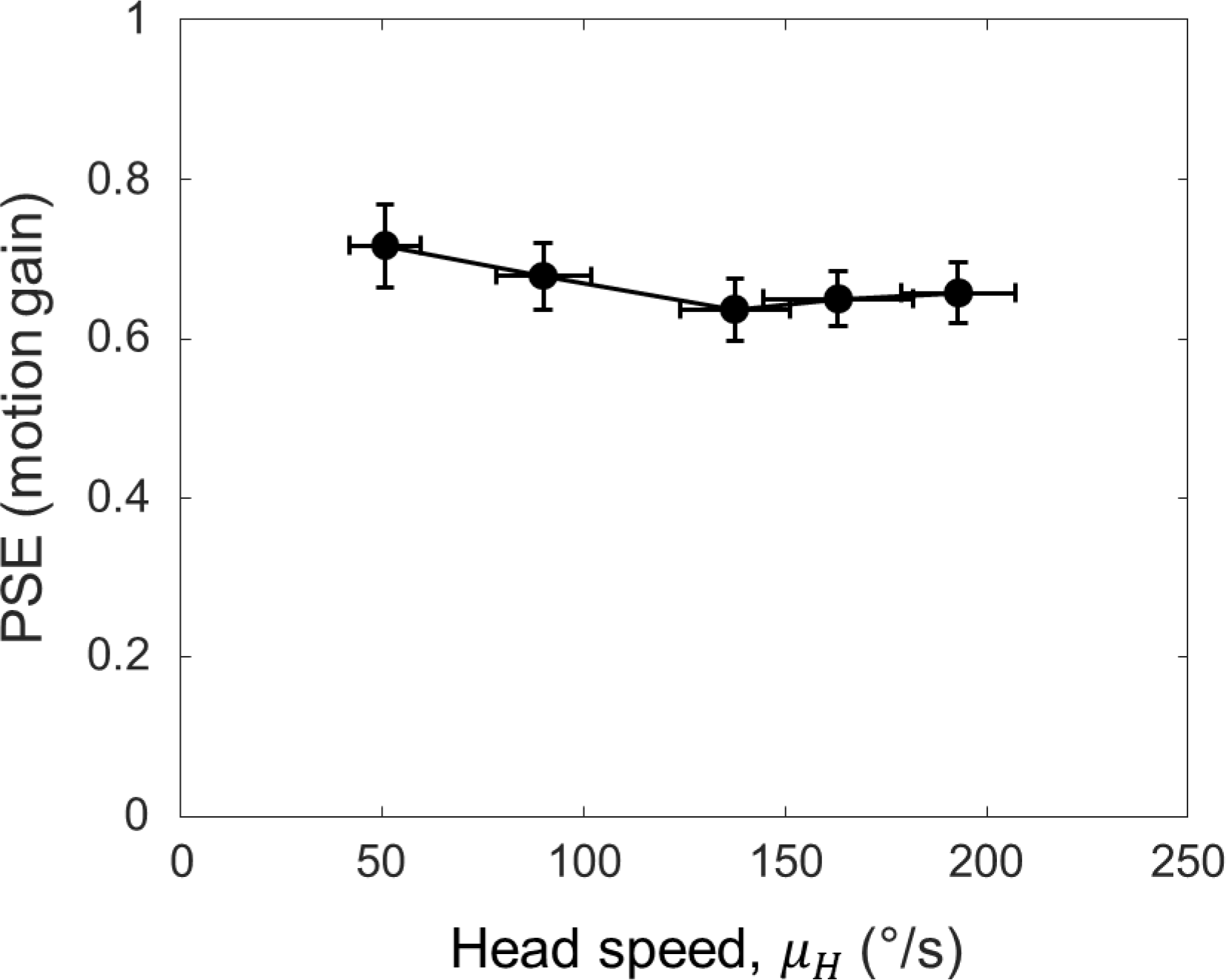
PSEs from Phase 1 as a function of head speed. Error bars are ±1SE.

For the auditory signal, precision did not vary with stimulus speed (F(4,45) = 0.154, p=.96). However, for the non-image signal, precision decreased with head speed, such that the standard deviation of the underlying signal distribution increased (F(4,45) = 6.035, p<0.001). Also evident is the fact that the auditory signal is less precise than the non-image signal over the range of stimulus speeds tested. Figure 7B plots the same data as Weber fractions i.e., standard deviation divided by head or stimulus speed. For both types of signal, precision adheres Weber’s law for medium to high speeds. Thus the Weber fractions are approximately constant over much of the range of speeds tested. At lower speeds, however, the two functions differ, with Weber fractions start to rise steeply for hearing. This rise is reminiscent of other studies of Weber’s law in the perception of auditory motion (Altman & Viskov, 1977). The same is not true for the non-image signal, where Weber’s law appears to hold reasonably well across all head speeds investigated. This finding is similar to previous work using passive stimulation of the vestibular system (Mallery et al., 2010).

Figure 8 plots the mean PSEs from Phase 1 as a function of mean head speed. They appear similar for all head speeds experienced (F(4,45) = 0.57, p = .69). Hence the same proportional reduction in image speed was needed to match the perceived motion in the head movement interval. This value was around 0.7, replicating the findings in Experiment 1. Over a wide range of head speeds, therefore, moving auditory stimuli appear slower during head movement, akin to the Aubert-Fleischl phenomenon in vision. However, unlike vision and pursuit eye movement (Freeman et al., 2010; Powell et al., 2016), the non-image signal appears more precise than the auditory image signal, which could have important implications for the interpretation of the bias shown in Figure 8. This point is taken up in more detail in the General Discussion.

In Experiment 1, we found non-image signal precision was higher using visual stimuli compared to auditory stimuli. To investigate further, we fit a regression line to the non-image precisions in Figure 7A using Deming’s technique, a procedure that is used when both X and Y values are dependent measures with error (see Harrison, Freeman & Sumner, 2015). The result is shown in Figure 9, together with the two non-image precision values found in Experiment 1. The regression analysis shows good agreement between Experiments 1 and 2 for auditory stimuli. The precision value from Experiment 1 (open circle) falls very close to the regression line determined by Experiment 2. At the same time, however, the analysis casts further doubt on whether Weber’s law can explain the better non-image precision found using visual stimuli (open triangle). If Weber’s law were to account for the discrepancy in precision, the head speeds in the visual condition of Experiment 1 would need to be halved in order to shift the open triangle horizontally onto the regression line. Reasons why modality might affect non-image precision are taken up in the Discussion.

**Figure 9.**
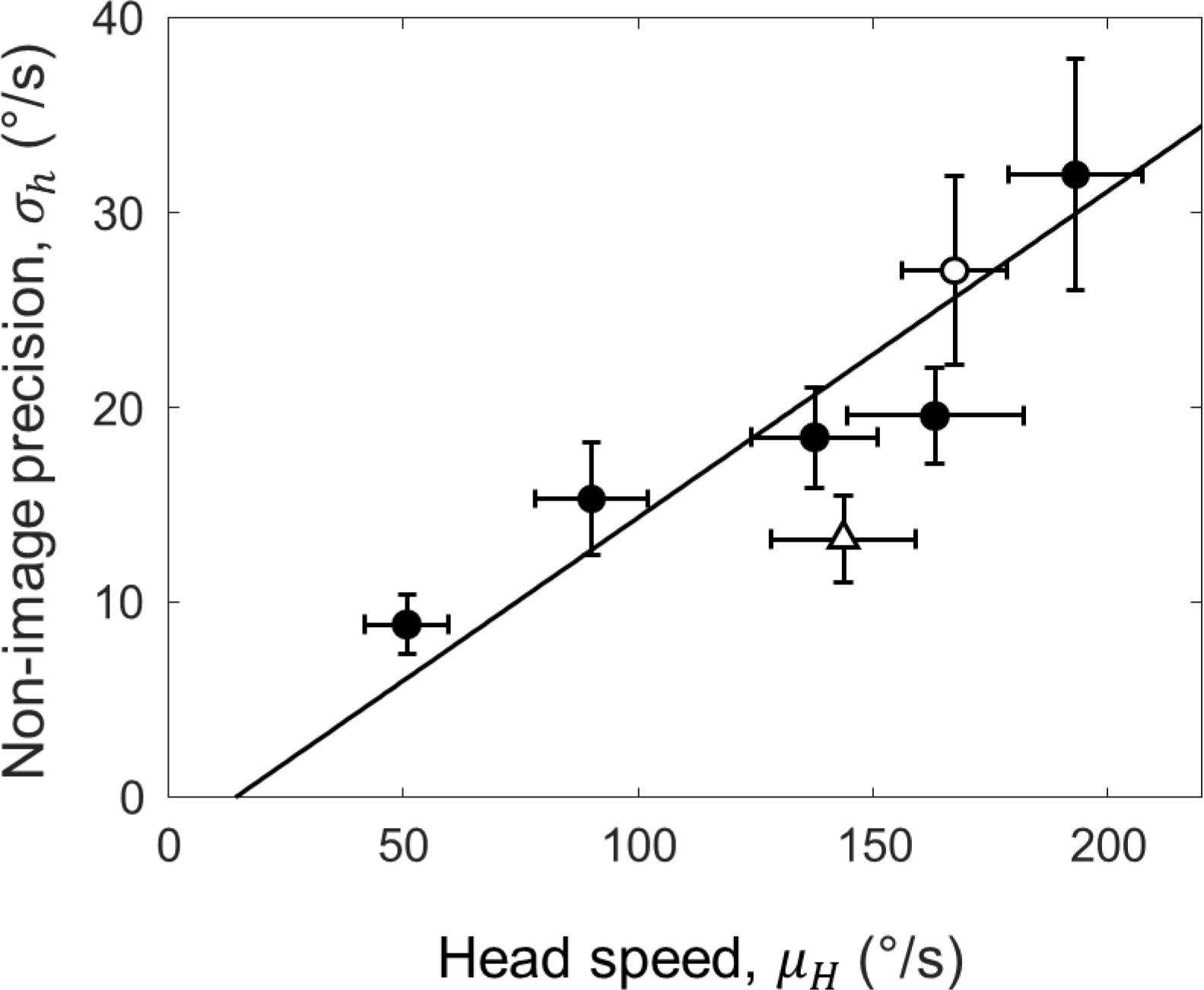
Deming regression (solid line) for the non-image signal precision data from Experiment 2 (filled circles). The open circle is the non-image signal precision from the auditory condition of Experiment 1, and the open triangle the visual condition. Error bars are ±1SE.

## General Discussion

We have proposed a novel technique for measuring the combined precision of the non-image signals that encode active self-movement (e.g. vestibular cues, proprioception and motor commands). Traditional psychophysical techniques are difficult to use in these situations because the self-movement is under participant control. Stimulation is therefore not repeatable across trials. The new technique relies on three factors: (1) linking image motion to self-movement using a motion gain parameter that can be manipulated in a consistent fashion across trials; (2) the generation of two psychometric functions limited by identical sources of noise, apart from the internal noise related to non-image signals encoding active self-movement; (3) a model that yields the internal noise sources, while controlling for the external noise created by self-movement as it varies across trials. The technique could easily be adapted for other examples of active self-movement, such as walking and active touch, and situations where non-image signals and image signals are experienced simultaneously (e.g. virtual reality).

### Assumptions of the model

The model assumes that eye movements made in all head-stationary test intervals are similar. If this were not the case, then the image noise in Phase 2 could be different from the noise in the test interval of Phase 1. For instance, if observers pursued the stimuli in the test interval of Phase 1, but not Phase 2, then motor cues related to the pursuit system would be present in the first phase but not the second. However, the eye movement recordings in Experiment 2 showed that fixation accuracy was similar in all test intervals, with the evidence suggesting that observers were able to keep their eyes stationary for a large percentage of the time. Fixation accuracy dipped for the head-moving interval, but most samples were still near the auditory target. Hence the eyes were not moving significantly for much of the time. While we did not measure eye movements in Experiment 1, it is unlikely we’d find fixation accuracy differed across test intervals in the two phases for vision and hearing. We say this for two reasons. First, we’d expect more variability in eye fixation to head-fixed auditory stimuli than visual stimuli, and yet fixation accuracy was good in Experiment 2. Second, unpublished results from our lab show that fixation accuracy is similar across phases for both visual and auditory stimuli.

The model assumes that internal noise is fixed. At first sight, this seems at loggerheads with the findings of Experiment 2, which show that the precision of the non-image signal declined as head speed increased. However, while the internal noises were fixed for each head-speed condition, they are a free parameter which could change across conditions. We view the fixed noise assumption as a reasonable approximation for the range of stimuli contributing to any single psychometric function. Indeed, the fixed noise assumption is implicit when fitting a single cumulative Gaussian to the data, unless stimulus values are logged.

### Measurement noise

The psychometric functions derived from Phase 1 and 2 are based on the same measurements of head rotation. This controls for any effect of measurement noise introduced by the head tracker because the (head-stationary) test intervals are based on the same set of recordings, presented in identical order. The effect on the precision of the image signal in the two phases is therefore the same, such that any influence is cancelled out.

This does not mean that all of our conclusions are immune to the effect of measurement noise. A case in point is the comparison carried out in Experiment 2 because measurement noise could differentially affect the estimates of auditory precision, as these estimates are based on one phase only. However, the effect of measurement noise is likely small. The standard deviation of position samples output by our head tracker is ∼0.55 deg. This is considerably lower than the positional noise needed to produce significant changes in speed discrimination thresholds previously reported in vision (Bentvelzen et al., 2009; Rideaux & Welchman, 2020). For instance, in the ‘high noise’ condition of Bentvelzen et al, positional noise was added to their LED system with a standard deviation of 7.4deg at an update rate of 25 Hz. This produced thresholds that doubled compared to baseline. If the stimuli used in Experiment 2 had been visual, we would expect thresholds to change around 7.4% (i.e. 0.55/7.4 x 100%). Bentvelzen et al used a two-interval technique, so this equates to a 5.25% change in the standard deviation of the underlying signal. This is a much smaller difference than found between the auditory image signal and non-image signal in Experiment 2. Moreover, spatial hearing is considerably less precise than vision, suggesting the effect would be even smaller still.

### Vision versus Hearing

Experiment 1 showed that non-image signal precision was lower when using visual stimuli in the head-stationary test intervals. We also found head speeds were slightly lower too, but the suggestion that Weber’s law might explain the difference in precision was not supported by the regression analysis of Experiment 2. This showed that non-image precision obtained with visual stimuli remained some way from the regression line despite the good agreement across experiments for auditory stimuli. Here we speculate that these differences stem from the need to convert perceptual signals into common units.

Suppose that the non-image signal is dominated by vestibular cues. Vestibular activity is based on acceleration, but as pointed out in the Introduction, vision prefers speed (Freeman et al., 2018; Reisbeck & Gegenfurtner, 1999) while hearing prefers displacement (Carlile & Best, 2002; Freeman et al., 2014). The motion signals therefore start in different units and must be transformed before they can be compared. One strategy is to integrate the vestibular cue once to get speed for vision, and twice to get displacement for hearing. Each transformation step adds noise. Therefore, non-image precision will be lower when auditory stimuli are used.

Alternatively, let’s suppose that motor signals (proprioception and motor commands) dominate the non-image signal. Freeman, Cucu & Smith (2019) showed that observers prefer speed versus displacement and duration cues when judging the motion of a pursued target, and that the cue being used was extra-retinal (i.e. motor commands and/or proprioception). Assuming the same is true for head rotation, the motor signals in our experiments start out in speed units. No transformation is therefore needed when comparing motor signals to vision because vision prefers speed anyhow, but one transformation step is needed to get the preferred cue of displacement for hearing. Once again, the precision measured using auditory stimuli would be lower because it needs ore transformation steps. The situation is more complex if the non-image signal consists of both vestibular and motor cues because they would need to be converted into common units before comparing with vision or hearing (known as cue promotion in the cue combination literature: Landy et al, 1995). Nevertheless, hearing will always need an additional transformation step compared to vision, meaning that the precision of the non-image signal being should decrease whenever auditory stimuli are used.

### Weber’s law

In Experiment 2, we found that Weber’s law described the precision of both image and non-image signals for medium to high speeds. However, at low speeds Weber fractions for the auditory image signal rose steeply, unlike those for non-image precision. Both findings echo previous reports in the literature. For auditory motion based on ITDs, Altman & Viskov (1977) found Weber fractions were roughly constant from around 60-140 deg/s but rose steeply at lower speeds. For vision, the same rise at slower speeds is found but matched by a similar rise at faster speeds (De Bruyn and Orban, 1986). For passive vestibular stimulation, the trend is similar to the Weber fractions we found for the non-image image signal, although further analysis by Mallery et al (2010) showed that a power law with an exponent around 0.4 is better description of the raw thresholds than the straight line predicted by Weber’s law. Similar behaviour has been reported for the variability in the vestibulo-ocular reflex, an eye movement controlled by the vestibular system (Nouri & Karmali, 2018). One implication is that the non-image signal we measured is dominated by the vestibular system. This assumes that the precision of motor signals behaves differently but we are unaware of any studies that have tried to measure the precision of motor signals on their own.

### Bayesian models of motion perception

In both Experiments 1 and 2 we found that perceived speed was lower when the head rotated. The bias was very consistent across modalities (Experiment 1) and stimulus speed (Experiment 2), adding to a large body of evidence showing that non-image signals based on eye rotation, head rotation, and hand/arm movement typically provide lower estimates of motion magnitude than signals encoding image motion in vision, hearing and touch (see below). On the face of it, the bias between non-image signals and image signals is puzzling because one might expect this type of constant error to be calibrated out by the perceptual system. One possible explanation is that the bias results from a Bayesian observer trying to optimise precision. According to the Bayesian hypothesis, the fact that early signals are noisy means that perception needs to infer the state of the world by combining imprecise measurements with prior expectations about the world state. The result is a posterior distribution that has greater precision than the original measurements, but not necessarily greater accuracy. As signals become noisier, the position of the posterior is increasingly pulled towards the prior distribution such that accuracy shifts. For motion, the claim is that the prior peaks at 0 because most objects are at rest (Weiss et al., 2002). Hence, as signals become noisier, speed estimates reduce.

The Bayesian framework has been used to explain why perceived visual speed slows at low contrast (Stocker & Simoncelli, 2006), why pursued objects appear slower (Freeman et al., 2010; Powell et al., 2016), why moving sounds appear slower when presented against background noise (Senna et al., 2015) and why tactile stimuli appear slower when made noisier or ‘pursued’ by an hand/arm movement (Moscatelli et al., 2019). It can also be used to account for individual differences in motion perception (Powell et al., 2016). Nevertheless, the overarching theory is not without its detractors (e.g. Hammett et al., 2007; Hassan & Hammett, 2015; Thompson et al., 2006). One simple test is to correlate measures of precision (e.g. thresholds) with bias – the Bayesian hypothesis predicts that as precision declines, perceived speed should slow. Many of the papers cited above show this to be case. However, there are a growing number of reports that this isn’t always true.Some recent studies in vision, hearing and vestibular research have shown changes in bias with little change in precision (Freeman et al., 2017; Freeman & Powell, 2022; Hassan & Hammett, 2015), and vice versa (Rideaux & Welchman, 2020). The findings of Experiment 2 add to these seemingly ‘non-Bayesian’ set of results. They show that auditory motion signals are less precise than non-image signals, even though the latter produce substantially lower estimates of motion magnitude.

## Conclusions

We have presented a novel technique for the measurement of the precision of non-image signals encoding active self-movement. We used head rotation as an example of self-movement, and showed that the precision measured was different when using auditory versus visual stimuli, which may be caused by the additional transform that must take place for comparison between non-image and auditory image signals. In agreement with current literature, we found that the non-image signal obeys Weber’s law over a wide range of stimulus speeds, unlike its image-based counterpart. We also found that the magnitude of perceived motion is reduced during head movement for both vision and hearing. This finding is difficult to explain within a Bayesian framework because image precision was not greater than non-image precision over the wide range of stimulus speeds investigated.

## Acknowledgements

The work here was supported by a Leverhulme Trust project grant (RPG-2018-151)

## APPENDIX

Phase 1 consists of a head-movement interval followed by an image-motion interval. In the first, there is no image motion as the object is spatially linked to the movement of the participant. Perceived motion therefore depends on a point estimate (*h*) of the non-image signal encoding head rotation. We assume that *h* is corrupted by fixed additive Gaussian noise across trials. Using N(*μ, σ*) to denote a normal distribution with mean *μ* and standard deviation *σ*, the non-image signal is therefore distributed as *h* = *μ*_*h*_ + N(0, *σ*_*h*_). The mean *μ*_*h*_ depends on the head movement magnitude (*H*), which we also assume is normally distributed across trials (see Figure 3B in the main text). Perceived motion in interval 1 (*M*_1_) is therefore:

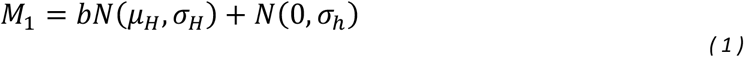

where *b* is a linear bias term that sets the gain of the head-movement signal relative to its input i.e. *h* = *bH*. Note that either speed or displacement could be used to characterise the distributions of head rotation and signals (to reiterate a point made in the main text, displacement and speed are perfectly correlated when manipulating motion gain because duration is fixed). The model is ambivalent. Swapping between speed and displacement changes the units but not the relative differences found for a chosen parameter across conditions.

In the second interval, image motion (*I*) moves as a fixed proportion (*g*) of the head movements recorded in interval 1: *I*(*t*) = *gH*(*t*). We refer to *g* as the ‘motion gain’. As the head and eyes are stationary, sensed movement depends on an image signal (*i*). Following similar logic to interval 1, the perceived motion in interval 2 is therefore:

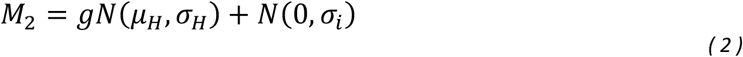

Note that Equation (3) assumes that the image signal is unbiased. Hence *b* in Equation 1 defines the *relative* bias between *h* and *i*, such that b < 1 means that the non-image signal registers a lower magnitude than the image signal. Following standard signal detection theory (e.g. Jones, 2016), we assume observers base their choice on an internal decision variable (*d*) that depends on the difference between the perceived motion in the two intervals:

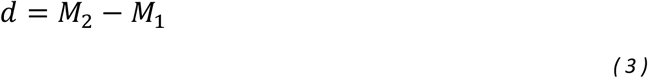

The choice ‘Interval 2 appears to move more’ corresponds to *d* > 0. From signal detection theory we define

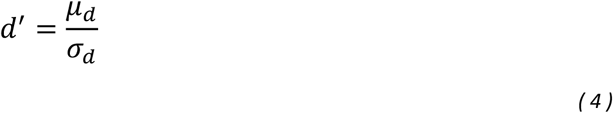

such that the probability of choosing Interval 2 is given by:

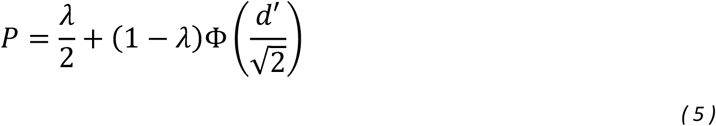

where *λ* is the lapse rate and Φ is the cumulative distribution function of the standard normal distribution.

Substituting (2) and (3) into (4):

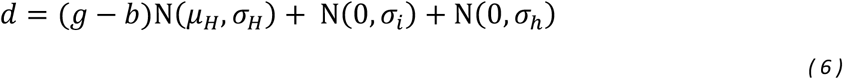

By inspection:

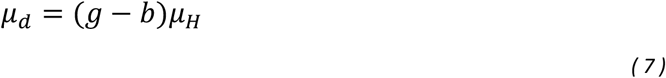

Note that the PSE occurs when *μ*_*d*_ = 0. At this point *g* = *b*; hence the relative bias between *h* and *i* can be read directly from the psychometric function. If the bias *b* < 1, then the PSE occurs when image motion is slower than head-movement. This is analogous to the Aubert-Fleischl phenomenon (Aubert, 1887; Fleischl, 1882), in which moving objects appear slower when pursued. Conversely, if *b* > 1, then image motion must be faster to achieve the PSE.

To obtain *σ*_*d*_, we sum the variances of the three distributions defined by equation (7) and take their square root:

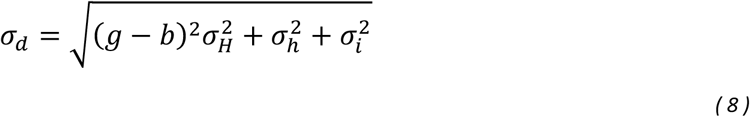

If the head movement did not vary across trials 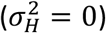, then the square root of the sum 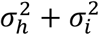 is the slope of the best-fitting cumulative Gaussian. The precision of the non-image signal (*σ*_*h*_) could then be obtained by measuring 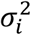 in Phase 2 and subtracting it from the sum. However, 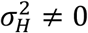. Variable head movements make the recovery of *σ*_*h*_ more complicated because they act as an external source of noise that varies with motion gain across the psychometric function.

**Figure A1:**
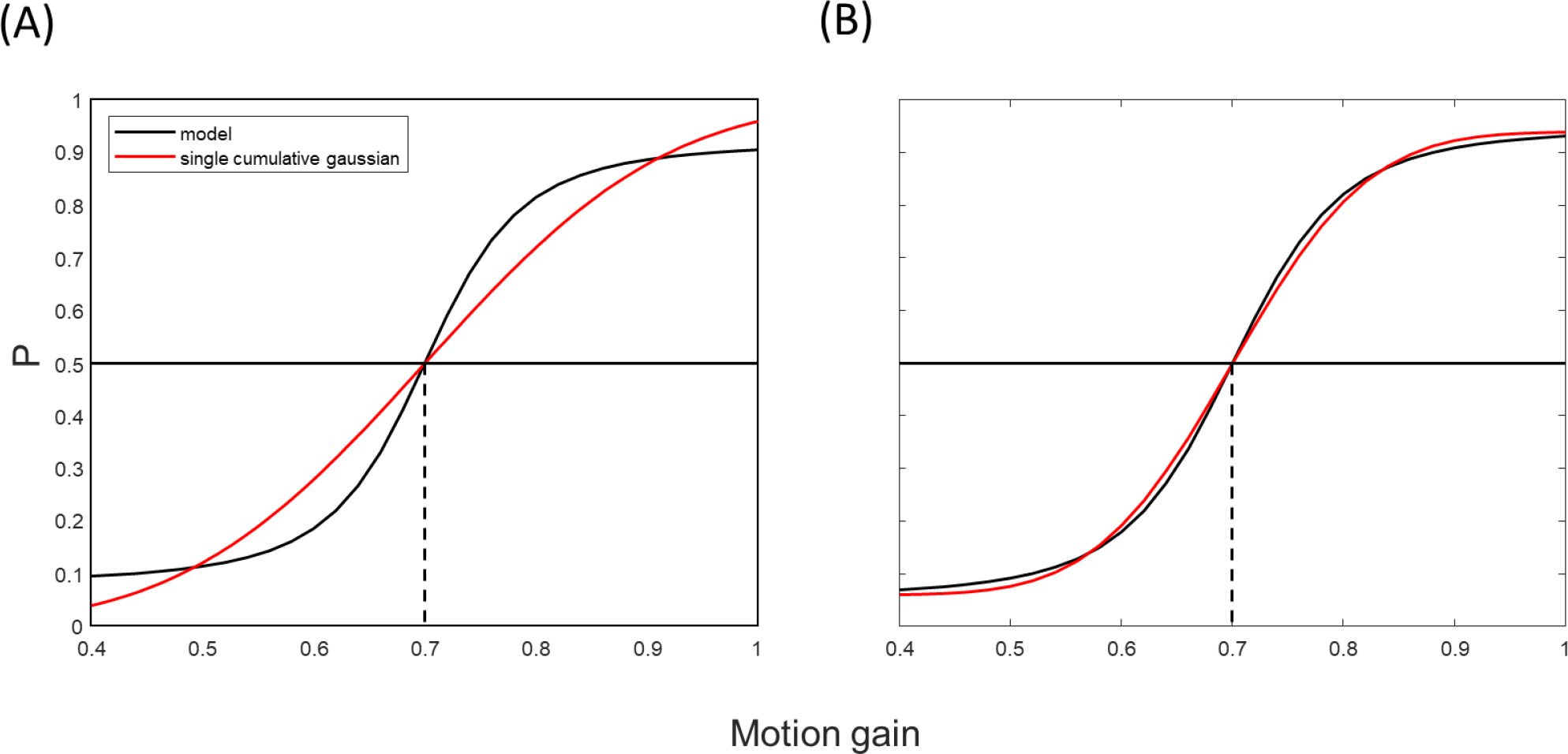
(A) The black curve shows a psychometric function based on gain-dependent noise with 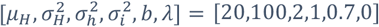. The red curve is the best fitting cumulative Gaussian as determined by the Palamedes toolbox, with lapse rate λ = 0. (B) Same curves but with lapse-rate λ = 0.06 for the gain-dependent noise psychometric function, and free to vary for the single cumulative Gaussian (λ ≤ 0.06, a standard constraint suggested by Wichman & Hill, 2001).

Figure A1 shows that fitting a single cumulative Gaussian is an approximation at best. The black curves show example psychometric functions based on the formulae above (see legend for parameter values) while the red curves show the best-fitting single cumulative Gaussian. The difference between the two panels is whether a lapse-rate is included or not. The external noise has two effects: (1) the asymptotes of the psychometric function move away from *P*= 0 and 1; (2) the slope becomes steeper and is not well fit by a single cumulative Gaussian. The degree to which the external noise causes substantial departures from the standard fit depends on the relationship between the values of 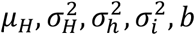 and whether lapse-rate is allowed to vary in the standard fit.

### Fitting procedure

We fit psychometric functions to our data based on the formulae above, using the measured head movements to estimate the mean and standard deviation of *H*. A matlab function for doing this can be found in the Supplementary Material (“fitSMmodel”). Phase 2 data were fit first, with 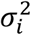 and *λ* free to vary, and *μ*_*H*_ and 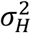 fixed, using the Gaussian distributions that we fit to the obtained head movement speeds (see Figure 3B in the main text). Phase 1 was then fit, with 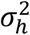, *b* and *λ* free to vary and 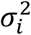, *μ*_*H*_ and 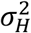 fixed. To avoid local minima in the fit, each parameter was cycled through a search space of 20 values and the best fit chosen. This yielded 20^n^ separate cycles of the fitting routine, where n is the number of free parameters which was different for the two phases.

We did not find much difference between fitting the new psychometric function and fitting a single cumulative Gaussian. One likely explanation for this similarity was that the head movements were relatively consistent 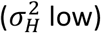 given the repetitive nature of the task. It may also be the case that including a constrained lapse-rate parameter soaked up a proportion of the asymptotic effect of the external noise. This can be seen by comparing Figure A1A (no lapse rate) with Figure A1B (constrained lapse rate <= 6%). The lapse rate mimics the asymptotic behaviour produced by the external noise.

### Region-of-interest for calculating head rotation speed

The analysis depends on mean and variance of the head movements made in interval 1 (see equations 8 and 9). The mean and variance were estimated from histograms of average speeds in the 3^rd^ sweep as described in the main text. To determine the region-of-interest (ROI), we compared the goodness-of-fit of psychometric functions from three ROIs: 20-80%, 20-60% or 40-60% of the sweep length. The psychometric functions were fit using MLE, so the appropriate measure of goodness-of-fit is the deviance (Wichman & Hill, 2001). Figure A2A shows that deviance in Experiment 1 did not change with the different ROIs used (the deviance has been averaged across conditions, phases and participants). The same was true for Experiment 2 (not shown). However, Figure A2B shows that an ROI of 20-80% produced a slower estimate of head movement speed than the other two ROIs, which was also more variable due to the inclusion of salient periods of acceleration and deceleration. Again the same was true for Experiment 2 (not shown). We therefore opted for an ROI of 20-60%.

**Figure A2:**
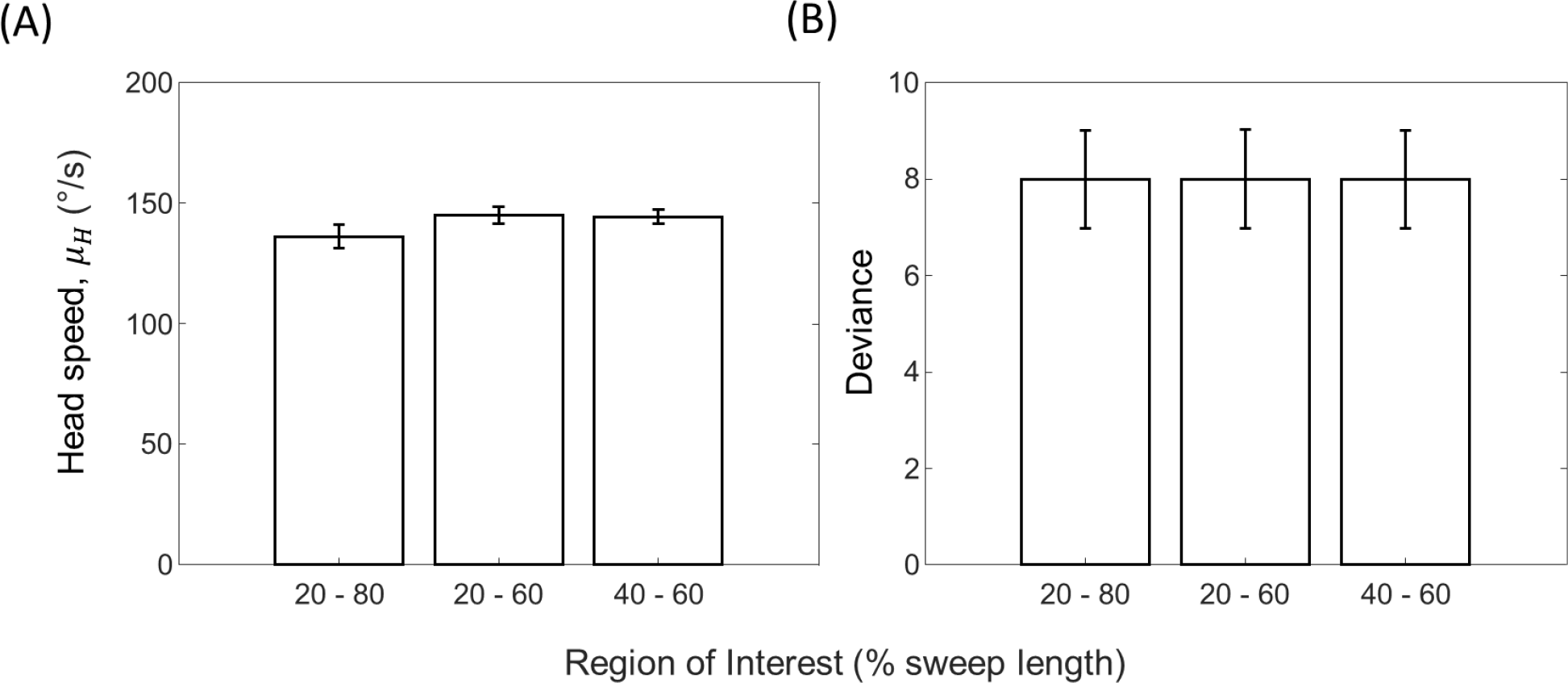
(A) Head speed and (B) model goodness-of-fit for each ROI used to analyse the head movements in Experiment 1. Error bars are ±1SE.

