## Supplementary material for "The precision of signals encoding active self-movement": matlab code (from another version of this preprint: version 1, 2023-09-22)

```
function [best_model, pse2, gof1, gof2] = fitSMmodel(gain1, gain2, P1, P2, pse1, n_per, H_mn,
H_var, max_lapse, error_type, n_search, is_plot)
% Inputs
%   gain1,gain2 = motion gains used in experiment for Phase 1 and 2
%   P1,P2 = probability choosing interval 2 moved more
%   pse1 = pse from Phase 1 used to set standard in Phase 2, in motion gain units
%   n_per = number of trials per motion gain level in pmf, assumed all the same
%   H_mn = mean measured head velocity
%   H_var = standard deviation
%   max_lapse = typically < 0.06, as per wichman & hill 2001
%   error_type = error to miniise on; least-squares 'ls' or 'mle'
%
% Outputs:
%   best_model = fitted model params:
%       (1) = bias as a proportion of head signal, latter in units set by H_* (= degree of
aubert, <1 classic, >1 reversed)
%       (2) = h_var, head signal variance, again in units set by H_*
%       (3) = i_var, image signal variance, ditto
%       (4) = lapse_rate for phase 1
%       (5) = lapse_rate for phase 2
%   pse2 = pse of best fitting pmf in phase 2, in motion gain units
%   gof1, gof2 = goodness-of-fit measures for Phase 1 and 2 [rms, deviance]
%
% Strategy:
% Fit Phase 2 data first with pse, i_var and lapse_rate free to vary
% Fit Phase 1 with bias, h_var and lapse_rate free to vary, but i_var now fixed by previous step.
%
% Avoid local minima using n_search to set granularity of search space per
% free variable, with ranges based on educated guesses as defined below
%
% Tom Freeman, c.June 2023

global max_lapse

% from fitpmf: when error_type = 'mle', this get empirical Ps equal to 0 or 1 away from floor or
% ceiling so log likelihood can be calculated without returning NaN
tweak = 0.001;
P1(P1==0) = tweak;
P1(P1==1) = P1(P1==1)-tweak;
P2(P2==0) = tweak;
P2(P2==1) = P2(P2==1)-tweak;

options = optimset('Display', 'off', 'FunValCheck', 'on');

try_var = logspace(0, 2, n_search); % either image or head signal
try_lapse = linspace(0, max_lapse, n_search);
try_bias = linspace(0.5, 1.5, n_search);

% fit phase 2 first to get i_var
err_1 = inf;
params_phase2 = [];
iii = 0; n = n_search^3; z = 4; zz = num2str(z); % merely for text output to command line to show
how far its got
mssg = 'phase 2: ';
fprintf('%10s',mssg)
for i = 1 : n_search
    for r = 1 : n_search
        for b = 1 : n_search
            iii=iii+1; fprintf(['%' zz 'i'],n-iii+1)
            start = [try_bias(b) try_var(i) try_lapse(r)];
            [current_best, ~, exitflag] = fminsearch(@fitfunc_phase2, start, options, gain2, P2,
n_per, pse1, H_mn, H_var, error_type);
            current_best = current_best.^2; % see fitfuncs
            err = error_metric(P2, n_per, pmf_phase2(current_best, gain2, pse1, H_mn, H_var),
error_type);
            if exitflag && (err < err_1)
                err_1 = err;
                params_phase2 = current_best;
            end
            for s = 1 : z, fprintf('\b'), end
        end
```

```
    end
end
for s = 1 : 10, fprintf('\b'), end
pse2 = params_phase2(1)*pse1; % i.e. convert into motion gain units, see pmf_phase2: this is for
output and plotting only

% Now fit phase 1 to get bias and h_var with i_var fixed by above
gof1 = NaN;
gof2 = NaN;
best_model = NaN;
mssg = 'phase 1: ';
fprintf('%10s',mssg)
if ~isempty(params_phase2)
    i_var = params_phase2(2);
    params_phase1 = [];
    err_1 = inf;
    iii = 0;
    for i = 1 : n_search
        for r = 1 : n_search
            for b = 1 : n_search
                iii=iii+1; fprintf(['%' zz 'i'],n-iii+1)
                start = [try_bias(b) try_var(i) try_lapse(r)];
                [current_best, ~, exitflag] = fminsearch(@fitfunc_phase1, start, options, gain1,
P1, n_per, H_mn, H_var, i_var, error_type);
                current_best = current_best.^2;
                err = error_metric(P1, n_per, pmf_phase1(current_best, gain1, H_mn, H_var,
i_var), error_type);
                if exitflag && (err < err_1)
                    err_1 = err;
                    params_phase1 = current_best;
                end
                for s = 1 : z, fprintf('\b'), end
            end
        end
    end
    for s = 1 : 10, fprintf('\b'), end

    gof1(1) = error_metric(P1, n_per, pmf_phase1(params_phase1, gain1, H_mn, H_var, i_var),
'rms');
    gof2(1) = error_metric(P2, n_per, pmf_phase2(params_phase2, gain2, pse1, H_mn, H_var),
'rms');
    gof1(2) = error_metric(P1, n_per, pmf_phase1(params_phase1, gain1, H_mn, H_var, i_var),
'dev');
    gof2(2) = error_metric(P2, n_per, pmf_phase2(params_phase2, gain2, pse1, H_mn, H_var),
'dev');

end

if is_plot
    sz = 30;
    x0 = min([gain1 gain2]);
    x1 = max([gain1 gain2]);
    savefig = gcf;
    figure(101), clf
    subplot(211), hold on
    x = linspace(x0, x1, 100);
    pse = params_phase1(1);
    if isnan(params_phase1)
       plot(gain1, P1, 'rx')
    else
        plot(x, pmf_phase1(params_phase1, x, H_mn, H_var, i_var), 'k-')
        plot(gain1, P1, 'k.','markersize',sz)
        line([x0 pse], [0.5 0.5], 'linestyle',':','color','k')
        line([pse pse], [0.5 0], 'linestyle',':','color','k')
    end
    title('Phase 1')
    ylabel('P')
    ylim([0 1])
    xlim([x0 x1])
    subplot(212), hold on
    if isnan(params_phase2)
```

```
        plot(gain2, P2, 'rx')
    else
        pp = pmf_phase2(params_phase2, x, pse1, H_mn, H_var);
        plot(x, pp, 'k-')
        plot(gain2, P2, 'k.','markersize',sz)
        line([x0 pse2], [0.5 0.5], 'linestyle',':','color','k')
        line([pse2 pse2], [0.5 0], 'linestyle',':','color','k')
    end
    title('Phase 2')
    xlabel('gain')
    ylabel('P')
    ylim([0 1])
    xlim([x0 x1])
    figure(savefig.Number)
end


% reordered output for backwards compatibility with older local analyses scripts
% pse2 returned separately as isn't really part of model
best_model = [params_phase1(1:2) i_var params_phase1(3) params_phase2(3)]; % bias, h_var, i_var,
lapse_rate1,lapse_rate2




function err = fitfunc_phase1(params, gain, P_data, n_per, H_mn, H_var, i_var, error_type)
global max_lapse
params = params.^2; % force values to be positive, but needs to square best_fit params outside to
return correct values
P_model = pmf_phase1(params, gain, H_mn, H_var, i_var);
err = error_metric(P_data, n_per, P_model, error_type);
if params(3)>max_lapse.^2+0.0001, err = inf; end % gets max_lapse = 0 working


function P = pmf_phase1(params, gain, H_mn, H_var, i_var)
bias = params(1);
h_var = params(2);
lapse_rate = params(3);
mn_d = (gain-bias)*H_mn;
sd_d = sqrt( (gain-bias).^2*H_var + h_var + i_var );
sd_d(sd_d==0) = 0.0001;% avoids /0 on next line
z = mn_d./sd_d;
P = lapse_rate/2 + (1-lapse_rate)*((1 + erf(z./sqrt(2)))/2);



function err = fitfunc_phase2(params, gain, P_data, n_per, pse1, H_mn, H_var, error_type)
global max_lapse
params = params.^2;
P_model = pmf_phase2(params, gain, pse1, H_mn, H_var);
err = error_metric(P_data, n_per, P_model, error_type);
if params(3)>max_lapse.^2+0.0001, err = inf; end % gets max_lapse = 0 working

function P = pmf_phase2(params, gain, pse1, H_mn, H_var)
alpha = params(1); % the pse for the pmf, as a proportion, see mn_d below; takes account of
sampling error, button bias etc
i_var = params(2);
lapse_rate = params(3);
mn_d = (gain-alpha*pse1)*H_mn;
sd_d = sqrt( (gain-pse1).^2*H_var + 2*i_var );
sd_d(sd_d==0) = 0.0001;% avoids /0 on next line
z = mn_d./sd_d;
P = lapse_rate/2 + (1-lapse_rate)*((1 + erf(z./sqrt(2)))/2);


% deviance calculation from psignifit, Wichman & Hill
function dev = deviance(y, n, p);
n = round(n);
r = round(n .* y);
w = n - r;
y = r ./ n;
```

```
residuals = 2 * (xlogy(r, y) + xlogy(w, 1-y) - xlogy(r, p) - xlogy(w, 1-p));
residuals(residuals < 0) = 0;
dev = sum(residuals, 2);
function a = xlogy(x, y)
k = (y==0);
y(k) = nan;
y(x==0 & k) = 1;
a = x.*log(y);
k = isnan(y);
a(k) = -sign(x(k)) * inf;

function err = error_metric(data, n, model, error_type)
% log likelihood
if strcmp(error_type, 'mle')
    err = sum( n.*data.*log(data./model) + n.*(1-data).*log( (1-data)./(1-model)) ); % from Klein
(2001)
% least-squares
elseif strcmp(error_type, 'ls')
    err = sum( (data-model).^2 )
elseif strcmp(error_type, 'rms')
    err = sqrt(mean((data-model).^2));
elseif strcmp(error_type, 'dev')
    err = deviance(data, n, model);
else
    error('**fitSMmodel: unknown error_type')
end
```
